# The developmental trajectory of ^1^H-MRS brain metabolites from childhood to adulthood

**DOI:** 10.1101/2023.10.05.560892

**Authors:** Alice R. Thomson, Hannah Hwa, Duanghathai Pasanta, Benjamin Hopwood, Helen J. Powell, Ross Lawrence, Zeus Garcia Tabuenca, Tomoki Arichi, Richard A. E. Edden, Xiaoqian Chai, Nicolaas A. Puts

## Abstract

Human brain development is ongoing throughout childhood, with for example myelination of nerve fibres and refinement of synaptic connections continuing until early adulthood. *^1^H-Magnetic Resonance Spectroscopy (*^1^H-MRS) can be used to quantify the concentrations of endogenous metabolites (e.g., glutamate and γ-aminobutyric acid (GABA)) in the human brain in vivo and so can provide valuable, tractable insight into the biochemical processes that support postnatal neurodevelopment. This can feasibly provide new insight into and aid management of neurodevelopmental disorders by providing chemical markers of atypical development. This study aims to characterize the normative developmental trajectory of various brain metabolites, as measured by ^1^H-MRS from a midline posterior parietal voxel. We find significant non-linear trajectories for GABA+, Glx, tNAA and tCr concentrations. Glx and GABA+ concentrations steeply decrease across childhood. tNAA concentrations are relatively stable in childhood but gradually decrease from early adulthood, while tCr concentrations increase from childhood to early adulthood. tCho was the only metabolite to have a strictly linear association with age. Trajectories likely reflect fundamental neurodevelopmental processes (including local circuit refinement) which occur from childhood to early adulthood and can be associated with cognitive development; we find GABA+ concentrations significantly positively correlate with recognition memory scores across post-natal development.

## 1. Introduction

There is increasing interest in developing measures of neurotypical brain development to understand the underlying biological processes and provide a reference for assessing atypical developmental trajectories. Magnetic resonance imaging (MRI) and histological studies have provided evidence of extensive postnatal human brain development, for example, there is a widespread increase in cerebral white matter volume and nerve fibre myelination spanning from birth to early adulthood (Lebel et al. 2008; Jernigan et al. 2011; Bethlehem at al., 2022). Concurrent with this is the refinement of regional and local brain circuitry as the initially excessive synaptic connections undergo activity dependent pruning (Changeux and Danchin 1976; Lebel et al. 2008; Jernigan et al. 2011). Accordingly, human cortical synaptic density peaks at 2-4 years of age before decreasing until early adulthood (Huttenlocher et al. 1979; Bourgeois et al. 1994). These changes in brain architecture correlate with various developmental changes in behaviour and cognition, including language, working and autobiographical memory (Hampson et al. 2006; Given-Wilson et al. 2018). The vast majority of work has however focused on structural changes in the brain’s architecture, with much less known about the accompanying changes in brain metabolism and neurotransmission.

*^1^H-Magnetic Resonance Spectroscopy (*^1^H-MRS) is a non-invasive MRI technique that can be used to quantify the concentrations of endogenous metabolites in the human brain in vivo (Puts and Edden 2012). ^1^H-MRS can thus provide valuable insights into the biochemical processes which likely underlie the observed structural brain development. Furthermore, understanding developmental changes in neurochemistry can feasibly provide much needed chemical biomarkers of atypical neurodevelopment, which crucially are likely to be more therapeutically tractable than existing structural indices (Blüml et al. 2013; Bethlehem at al. 2022).

^1^H-MRS commonly focuses on the brain’s high-concentration metabolites; N-acetyl aspartate (NAA), total choline (tCho) and total creatine (tCr). These molecules can be found in both neurons and glial cells (Rae et al. 2014). NAA is synthesised by N-acetyltransferase 8-like (NAT8L) in the mitochondria and plays a key role in metabolic pathways including oxidative phosphorylation (Patel and Clark 1979; Rae et al. 2014; Siddiqui et al. 2016). Choline is seldom found in a ‘free’ form, but within the choline-containing compounds phosphocholine, phosphatidylcholine and glycerophosphocholine, which together make up the composite MRS signal referred to as tCho (Miller et al. 1996; Rae et al. 2014). These molecules are essential for cell membrane integrity and signalling, as well as acetylcholine synthesis (within cholinergic neurons) and are found in myelin sheath (Rae et al. 2014). The ^1^H-MRS tCr signal is composed of creatine and its phosphorylated form phosphocreatine, formed when ATP reacts with creatine (a reaction catalysed by creatine kinase). This reaction is reversible and can more readily release ATP compared to oxidative phosphorylation, meaning phosphocreatine acts as a fast energy store (Walliman et al. 1992; Rae et al. 2014). Understanding the maturational trajectory of these metabolites can thus give insights into the neurochemical integrity of the brain with more mechanistic insights compared to structural imaging alone.

In addition, specifically tailored ^1^H-MRS sequences can be used to estimate the concentrations of the primary inhibitory and excitatory neurotransmitters of the adult human brain, γ-aminobutyric acid (GABA) and glutamate (Glu), along with their precursor glutamine (Gln) (Puts and Edden 2012; Mullins et al. 2014; Harris et al. 2017). GABA, Glu, and Gln, have fundamental roles in synaptic neurotransmission and metabolism, including regulating neuronal connectivity, and myelination (Madhavarao et al. 2005; Ghisleni et al. 2015; Serrano-Regal et al. 2020). Notably during development, GABA and glutamate are not ‘inhibitory’ or ‘excitatory’ per se, with GABA signalling known to evoke neuronal depolarisation (excitation) early in postnatal development in many species including mammals (Dzhala et al. 2005; Kirmse et al. 2015).

Previous attempts to characterise the neurodevelopmental dynamics of these key brain metabolites have considered narrow adult only (≥18) or child only (<18) cohorts (Chang et al. 1996; Saunders et al. 1999; Pfefferbaun et al. 1999; Angelie et al. 2001; Kaiser et al. 2005; Sailasuta et al. 2008; Reyngoudt et al. 2012; Marsman et al. 2013; Gao et al. 2013; Blüml et al. 2013; Hädel et al. 2013; Silveri et al. 2013; Ding et al. 2016; Eylers et al. 2016; Puts et al. 2017; Schmitz et al. 2018; Lind et al. 2020; Saleh et al. 2020; Hupfeld et al. 2021), and/or have focused on very few metabolites (Ghisleni et al. 2015; Volk et al. 2019; Shimizu et al. 2017). Combined with small sample sizes, variation in brain region, and ^1^H-MRS quantification methods, this has led to highly conflicting results. As such, deciphering individual metabolite developmental trajectories that are concurrent with the aforementioned known structural changes remain a challenge (Porges et al. 2021). Furthermore, most studies to date have used linear modelling approaches for metabolite trajectories, which may fail to capture subtle differences across development and thus do not truly reflect the rich and complex changes that likely occur during maturation (Lebel et al. 2011; Porges et al. 2021).

Accordingly, here we **aim** to characterize the developmental trajectory of six brain metabolites (GABA, Glu, Gln, NAA, tCho and tCr), as measured by edited ^1^H-MRS from a posterior parietal cortex voxel (PPC) in a large typically developing sample (117 participants) from childhood to early adulthood (5 – 35 years old). As the PPC is associated with episodic memory formation and retrieval (Wagner et al. 2005; Vincent et al. 2006; Ciaramelli et al. 2008; Cabeza et al. 2008; Rugg and King 2018; Neri et al. 2021), we also aim to explore whether neurodevelopmental differences in ^1^H-MRS trajectories are associated with development of cognitive function, by observing if GABA/Glu concentrations in particular associate with recognition memory scores.

## 2. Methods

### Participants

117 participants participated in in vivo MRI imaging and cognitive testing. All participants consented to participation in the study through local Kennedy Krieger Institute and Johns Hopkins University IRB procedures. Children assented and their parents consented to participation. All participants were native English speakers, right-handed, had normal or corrected-to-normal vision, with no history of psychiatric, neurological, or developmental disorders. All participants had IQ > 85. Participant demographic data were collected at the time of scan. 31 MRS datasets were excluded due to poor data quality (10 during data processing, and 21 during quality data control. This was mostly due to motion artefacts; see quality control section below). As such our analysis uses data from 86 participants between 5 and 35 years of age (40 females, 46 males, mean age = 16.45 years, SD = 7.24). The range of participant IQ scores was 85 – 138 (median IQ = 118). Median household income of participants was > $100k (range: <$35K – >$100K).

### Memory task

Participants completed a source memory encoding task in the MRI after the MRS scan and were given a memory test after the MRI session. Encoding stimuli consisted of 4 blocks of 40 colour images of commonly known, visually distinct objects overlaid on top of one of two backgrounds (beach or forest). Object images fell under one of seven categories (animal, clothing, fruit, vegetable, toy, tool, instrument). For each trial, participants answered either one of the two questions: “Do you like this object or dislike/not care about it?”, indicated by a smiling cartoon face and a neutral cartoon face, or “Is this a living or not living object”, indicated by a leaf and a leaf with a red “X” through it. The background and encoding questions were randomly assigned to each object, ensuring that each category had an equal distribution of the four background and question combinations. Each image was shown for 3 seconds (s), followed by a fixation screen, consisting of a white “+” symbol on a black background, for 1 to 9 s.

The memory test consisted of three blocks of 80 images either from the encoding task or new images from the same categories, totaling 240 images (80 new, 160 previously seen during the encoding task). Images were displayed in pseudorandom order, with no more than three consecutive images from either the new image set or from the encoding activity. For each image, participants were asked to first determine whether they remembered seeing the object during the encoding activity and also remember specific details (e.g., what the image looked like on the screen, what they were thinking at the time, etc.), did not remember seeing the object, or thought the object was familiar but could not recall additional details (denoted by the options “Remember,” “New,” and “Familiar” respectively; Gardiner, 1988). If the participant chose either remember or familiar, they were then asked to answer which of the two backgrounds the object was shown with, followed by which of the two question prompts (living/ non-living, or like/do not like) the object was shown with. The test was self-paced in which the next question was presented as soon as the participant pressed a response key.

The trials were categorized based on the answers provided during the testing task into “Hits” (old objects correctly identified as remember or familiar), “Miss” (old objects identified as “New”), “False Alarm” (new objects falsely identified as remember or familiar), and “Correct Rejection” (new objects identified as “New”). Hits were further divided into remember and familiar based upon the participants’ answer. Remember and familiar trials were further divided into those with correct source information (background image correct, encoding question correct, or both sources correct). Recognition memory accuracy was calculated by subtracting the percentage of false alarms from the percentage of hits (%Hits–%FA).

### MRI/^1^H-MRS

MRI and ^1^H-MRS was performed on the Philips 3 Tesla Achieva scanner (Best, NL) at the F.M. Kirby Research Centre for Functional Brain Imaging at the Kennedy Krieger institute in Baltimore, USA. For all scans, a 32-channel head coil was used for receiver, and the body coil for transmit. Prior to MRS, a high-resolution (1 mm^3^ isotropic) T1-weighted MP-RAGE anatomical image was acquired for voxel placement and tissue segmentation.

MRS data were acquired from a 27 ml voxel (3 x 3 x 3 cm^3^) placed over the posterior parietal cortex, centred on the midline (see **Figure 1A**). MRS was performed using GABA selective MEshcher-Garwood Point RESolved Spectroscopy (MEGA-PRESS; Mescher et al. 1998) with the following parameters: 320 transients (160 ON and 160 OFF), 2048 data points, TE/TR 68/2000ms with editing pulses placed at 1.9ppm in the Edit-ON acquisitions and 7.46 in the Edit-OFF acquisitions and VAPOR water suppression. An interleaved unsuppressed water reference (Edden et al. 2016) with same parameters for water supressed scans was used (16 averages) to mitigate scanner drift and for subsequent eddy-current and phase corrections, and metabolite quantification.

**Figure 1.**
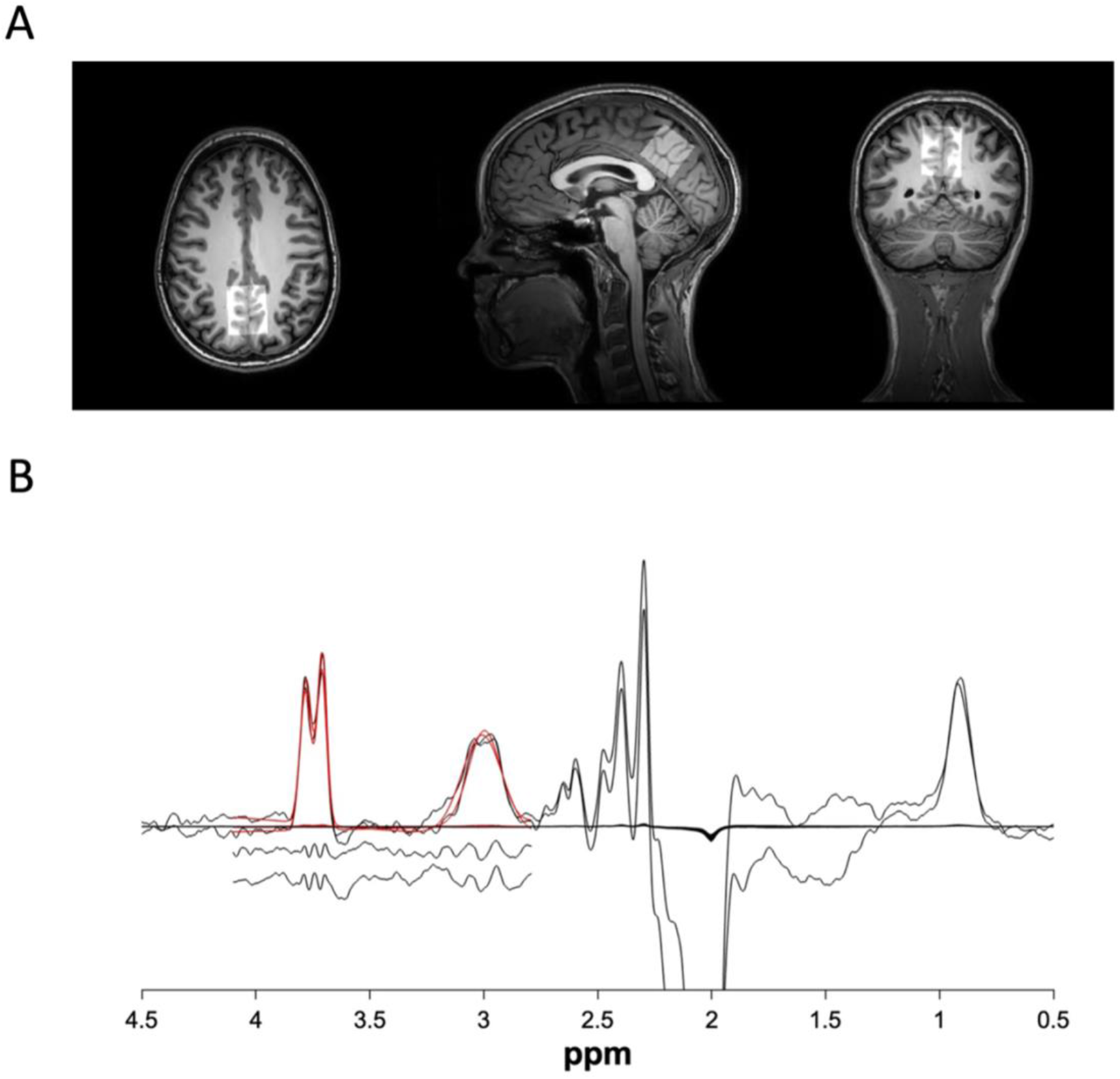
(A) Example voxel placement (B) average difference (ON – OFF) spectra (averaged across all participants).

### Data processing

Raw MRS data were processed using Osprey (Version 2.4.0; Oeltzschner et al. 2020), an automated software for MRS analysis based in Matlab (2022a, The Mathwors, Natick MA). Data from 10 participants could not be processed due to large motion artefacts identified in the frequency trace and poor spectral quality where no edited peaks could be identified. We used the following Osprey processing steps: first, raw data were eddy current-corrected using the water reference followed by frequency and phase correction using robust spectral registration (Near et al. 2015; Mikkelsen et al. 2020; Oeltzschner et al. 2020), Fourier transformation and water signal removal. Spectra were then fit between 0.5 and 4 ppm using baseline correction. Edit-ON and edit-OFF spectra averages were subtracted to resolve GABA (at 3.02 ppm) and Glx (at 3.56 ppm) from the overlaying metabolites (creatine, phosphocreatine, NAA), which are subtracted out in the difference spectrum. An averaged ON – OFF (difference) spectrum is shown in **Figure 1B**. GABA signals are known to be contaminated with macromolecules that are also affected by the GABA-selective editing pulse (for more information, see Mullins et al. 2014) and thus are reported as GABA+ (GABA + macromolecules). Average metabolite spectra were modelled using a TE-specific simulated basis set and a flexible spline baseline based on MRS vendor, pulse duration and scan sequence parameters (Oeltzschner et al. 2020). Linear-combination modelling (Provencher et al. 2001) was performed for 19 metabolites (ascorbic acid, aspartic acid, total Creatine, creatine methylene, GABA, glycerophosphocholine, glutathione, glutamine, glutamate, myo-inositol, lactate, total N-acetylaspartate, n-acetylaspartylglutamate, total choline, phosphocholine, phosphocreatine, phosphatidylethanolamine, scyllo-inositol, taurine) and 8 macromolecules (MM09, MM12, MM14, MM17, MM20, Lip09, Lip13, Lip20).

Here, we focus on GABA+, Glu, Gln and Glx (see below) as well as tNAA, tCr, and tCho. Due to the overlapping peaks of NAA and N-Acetylaspartylglutamic acid (NAAG) in the chemical shift spectra, their combined concentrations are denoted tNAA (total NAA). Glutamate and glutamine also overlap in the chemical shift spectra (at 3T) and so Glx is typically used to denote their combined signal (glutamate + glutamine). While we report Glx, Osprey uses fitting to additionally produce glutamate and glutamine only estimates. We report both Glx and Glu for transparency. Metabolite concentrations were estimated relative to the unsuppressed water signal in institutional units (i.u.) and as a ratio relative to total creatine (e.g., tCho/cr). GABA+, Glx and Glu were quantified in the subtracted edit ON-OFF spectra, while tNAA, tCho, tCr were quantified in the edit OFF spectra. The Osprey co-registration module (via SPM version 12) was used to register MRS to the T1-weighted images acquired at the scan, before segmenting the voxel volume into gray matter (GM), white matter (WM) and cerebrospinal fluid (CSF). Outputs were visually inspected to ensure accurate localization of the MRS voxel. Segmented T1 images were used to obtain tissue composition corrected (GM, WM and CSF) water-scaled estimates of metabolite concentrations (i.u.), whereby concentrations are scaled according to the assumption that metabolite concentrations in CSF are negligible (Gasparovic et al. 2006; Harris et al., 2015). Metabolite T1 and T2 relaxation effects were also accounted for (tissue water and metabolite; Gasparovic et al. 2006, Puts et al. 2013, Oeltzschner et al. 2020). Finally, ‘alpha correction’ of GABA+ concentrations was performed in Osprey, with the assumption that GABA+ concentration is two times larger in GM compared to WM (Jensen et al. 2005, Harris et al. 2015).

### Spectral artifacts and quality control

Spectra and Osprey quality metrics (SNR, FWHM, residual water, frequency shift, fit residuals) were visually inspected by an experienced MRS data user (NP) blind to participant age, and datasets with significant artifacts and/or poor quality (> 10% greater than average SNR & fit residuals) were immediately excluded (N = 21). Box plot inspection of metabolite values ensured no extremely significant QM outliers remained. Data (N = 86) are reported based on consensus guidelines (Lin et al. 2021). Quality metrics were added as co-variates to statistical models.

### Statistical analysis

Statistical analysis of data was performed on RStudio (2022.07.2). Data were analysed both across all participants with age as a continuous variable, and categorically by age group (**Table 1**). We are primarily interested in the developmental trajectory, but as many existing studies focus on specific age groups (childhood (5 - 12), adolescence (13 - 18), adulthood (18+)) we were keen to also establish differences between age categories to aid interpretation with respect to the literature. Data were first tested for normality using the Wilks-Shapiro test and tested for equal variance (of age categorical groups) using Levene’s test, confirming that parametric tests were suitable. Data are reported as mean (SD) unless stated otherwise.

**Table 1.** Participant demographics by age group.

| Age group | N (females) | Mean age (SD) | Mean IQ (SD) |
| --- | --- | --- | --- |
| Child ($5 \leq \text{age} < 12$) | 31 (15) | 9.86 (1.66) | 112.03 (13.26) |
| Adolescent ($12 \leq \text{age} < 18$) | 22 (10) | 14.44 (1.48) | 112.82 (11.48) |
| Adult ($\text{age} \geq 18$). | 33 (15) | 29.98 (5.63) | 119.88 (9.61) |
| All participants | 86 (40) | 16.45 (7.24) | 115.24 (11.96) |
| p value |  |  | 0.0157 * |

Note: p value indicates significant demographic difference between age groups (ANCOVA).

#### Group comparisons

Analysis of covariance tests (ANCOVA) were used to determine the impact of age on quality MRS metrics, demographic information, voxel tissue fractions and metabolite concentrations. Tukey HSD was used for post-hoc testing accounting for multiple comparisons and unequal groups sizes. IQ, sex and fit residuals were subsequently added to ANCOVAs as covariates. Sex had no significant effect for any metabolites (excluding tCho and Gln) and thus males and females were pooled for further analysis. For tCho and Gln, sex effects where further explored using Wilcoxon Signed Rank Tests with Bonferroni multiple comparison corrections. Correlation coefficients were calculated to determine the association between voxel tissue fractions and age using the *cor* function in R, as well as the correlations between metabolite concentrations and age for each predefined age group (**Table 1**).

#### Modelling metabolite trajectories

Both linear and non-linear regressions were used to estimate metabolite trajectories. For linear regressions we used the *lm* function in R, with metabolite as outcome and age as the predictor. Sex, IQ and fit residuals were successively added to the linear model and the resulting R squared was compared to isolate the most appropriate linear model, this being: *metabolite ∼ sex + fit residuals + IQ + age*. The likelihood ratio test was used to check the full model (IQ, sex, fit residuals, and age) against nested models to further confirm appropriateness. For tCho and Gln concentrations, exploratory linear regressions were performed for sex-separated data due to significant sex interactions.

For non-linear models we use Generalised Additive Models (GAM; Hastie and Tibshirani, 1990; Hastie 1992) using the ‘*gam’* function from the *mgcv* package in R. GAM is a generalised addictive modelling approach whereby the impact of the predictor variables on the dependent variable is modelled through a set number of flexible spline functions which can be linear or non-linear. Each spline making up the GAM has its own unique coefficient and splines are added together to produce the final model, from which we obtain several inferential parameters (R^2^, F statistic and p values). Neurometabolite concentrations were modelled using smooth (Gaussian process) functions of age, IQ and fit residuals, while sex was added as a categorical predictor. The Restricted Maximum Likelihood method (REML) was used to select the Gaussian process smoothing parameters for each predictor. *‘gam.check’* function was used to select a suitable number of smooth functions for each predictor (residuals should be randomly distributed). For GAM, the effective degrees of freedom (*edf*) for each smooth function (for each predictor) are assessed, these represent the complexity of the smoothing. Edf’s of 1 indicate a linear relationship between the dependant variable and predictor, while edfs of 2 indicate a quadratic relationship; greater *edf* values as such indicate a greater degree of non-linearity. Reference degrees of freedom and the F statistic were used to assess the significance of predictor smooth functions (significance meaning greater certainty of the shape of the smooth). Given the sex effects, sex-specific trajectories were plotted for Gln and tCho.

#### Correlational analysis

To measure correlations between metabolites (in order to observe the interaction between their concentrations across development), we used Pearson correlation coefficients for every metabolite pair before plotting onto a correlation matrix, this is reported for all metabolites in the supplementary materials. We did this per age category to assess whether metabolite cross-correlations qualitatively changed with age. We were particularly interested in the interaction between measures of brain excitation and inhibition (Glx/Glu/GABA+) given their known relationship to brain function.

Pearson correlation coefficients were also calculated between age and E/I ratios (estimated and creatine-ratio concentrations of Glx/Glu were divided by GABA+), and Glx/Glu/GABA+ concentrations and memory scores (recognition). The linear regression (*recognition memory ∼ age + GABA+)* was used to account for (and regress out) the effect of age to better explore the relationship between recognition memory and GABA+ concentrations.

## 3. Results

Participant demographics are reported in **Table 1**. Age group had a significant main effect on IQ (p < 0.05), with IQ significantly greater in adults compared to children and adolescents.

In line with consensus reporting standards (MRSinMRS; Lin et al. 2021, MRS-Q; Peak et al. 2020), we report data quality metrics (QM) to assess the quality of the MRS data and fit (**Table 2**). QM measures across subgroups met standard metrics confirming high quality spectra in the retained data (Mikkelsen et al. 2017).

**Table 2.**
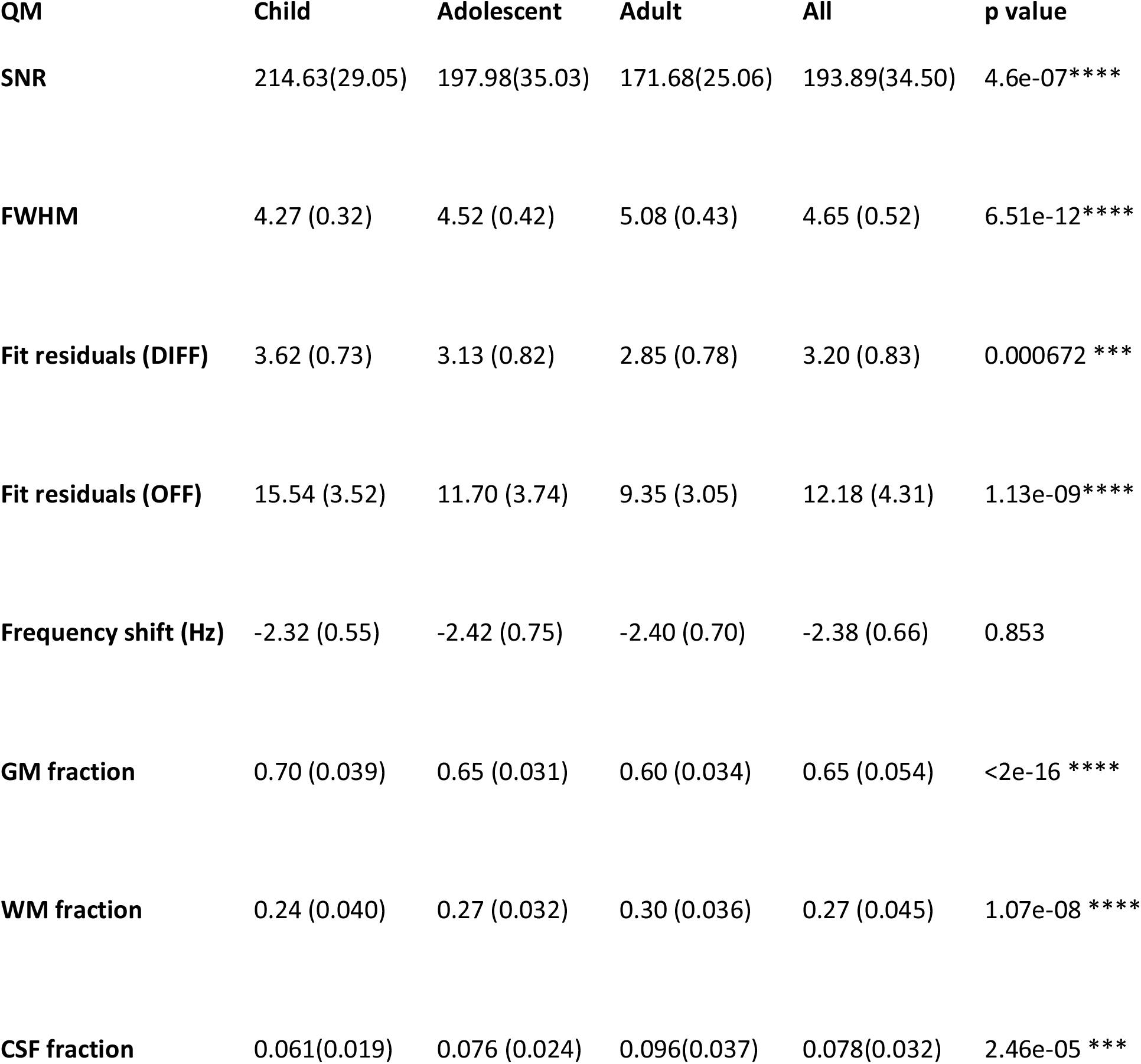
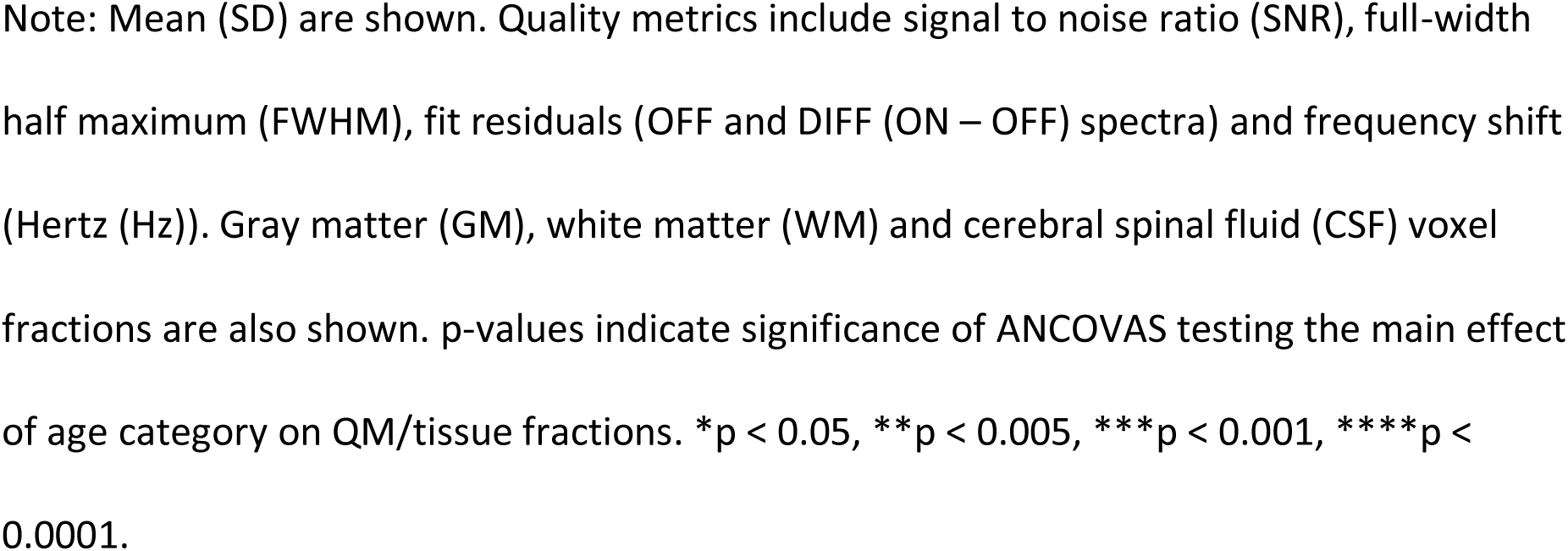
Quality of MRS data by age group.

| **QM** | **Child** | **Adolescent** | **Adult** | **All** | **p value** |
| --- | --- | --- | --- | --- | --- |
| **SNR** | 214.63(29.05) | 197.98(35.03) | 171.68(25.06) | 193.89(34.50) | 4.6e-07**** |
| **FWHM** | 4.27 (0.32) | 4.52 (0.42) | 5.08 (0.43) | 4.65 (0.52) | 6.51e-12**** |
| **Fit residuals (DIFF)** | 3.62 (0.73) | 3.13 (0.82) | 2.85 (0.78) | 3.20 (0.83) | 0.000672 *** |
| **Fit residuals (OFF)** | 15.54 (3.52) | 11.70 (3.74) | 9.35 (3.05) | 12.18 (4.31) | 1.13e-09**** |
| **Frequency shift (Hz)** | -2.32 (0.55) | -2.42 (0.75) | -2.40 (0.70) | -2.38 (0.66) | 0.853 |
| **GM fraction** | 0.70 (0.039) | 0.65 (0.031) | 0.60 (0.034) | 0.65 (0.054) | <2e-16 **** |
| **WM fraction** | 0.24 (0.040) | 0.27 (0.032) | 0.30 (0.036) | 0.27 (0.045) | 1.07e-08 **** |
| **CSF fraction** | 0.061(0.019) | 0.076 (0.024) | 0.096(0.037) | 0.078(0.032) | 2.46e-05 *** |

Note: Mean (SD) are shown. Quality metrics include signal to noise ratio (SNR), full-width half maximum (FWHM), fit residuals (OFF and DIFF (ON – OFF) spectra) and frequency shift (Hertz (Hz)). Gray matter (GM), white matter (WM) and cerebral spinal fluid (CSF) voxel fractions are also shown. p-values indicate significance of ANCOVAS testing the main effect of age category on QM/tissue fractions. \* $p < 0.05$ , \*\* $p < 0.005$ , \*\*\* $p < 0.001$ , \*\*\*\* $p < 0.0001$ .

Age group differences were observed for SNR, FWHM and spectral fit residuals. Tukey post hoc analysis shown in **Figure 2** found that SNR was significantly lower in the adult data compared to child (171.68 (25.06) versus 214.63 (29.05); p < 0.0001) and adolescent data (171.68 (25.06) versus 197.98 (34.50); p < 0.005), however SNR is still considered to be high for all age groups (Wilson et al. 2012). The adult data had significantly greater FWHM than child data (5.08 (0.43) versus 4.52 (0.42); p < 0.0001) and adolescent data (5.08 (0.43) versus 4.27 (0.32); p < 0.005), however FWHM is still considered small for each age group and represents a good shim (Wilson et al. 2012). Finally, spectra fit residuals of child data were significantly greater than that of adult data (3.62 versus 2.85; p < 0.005). We explore this in the discussion below.

**Figure 2.**
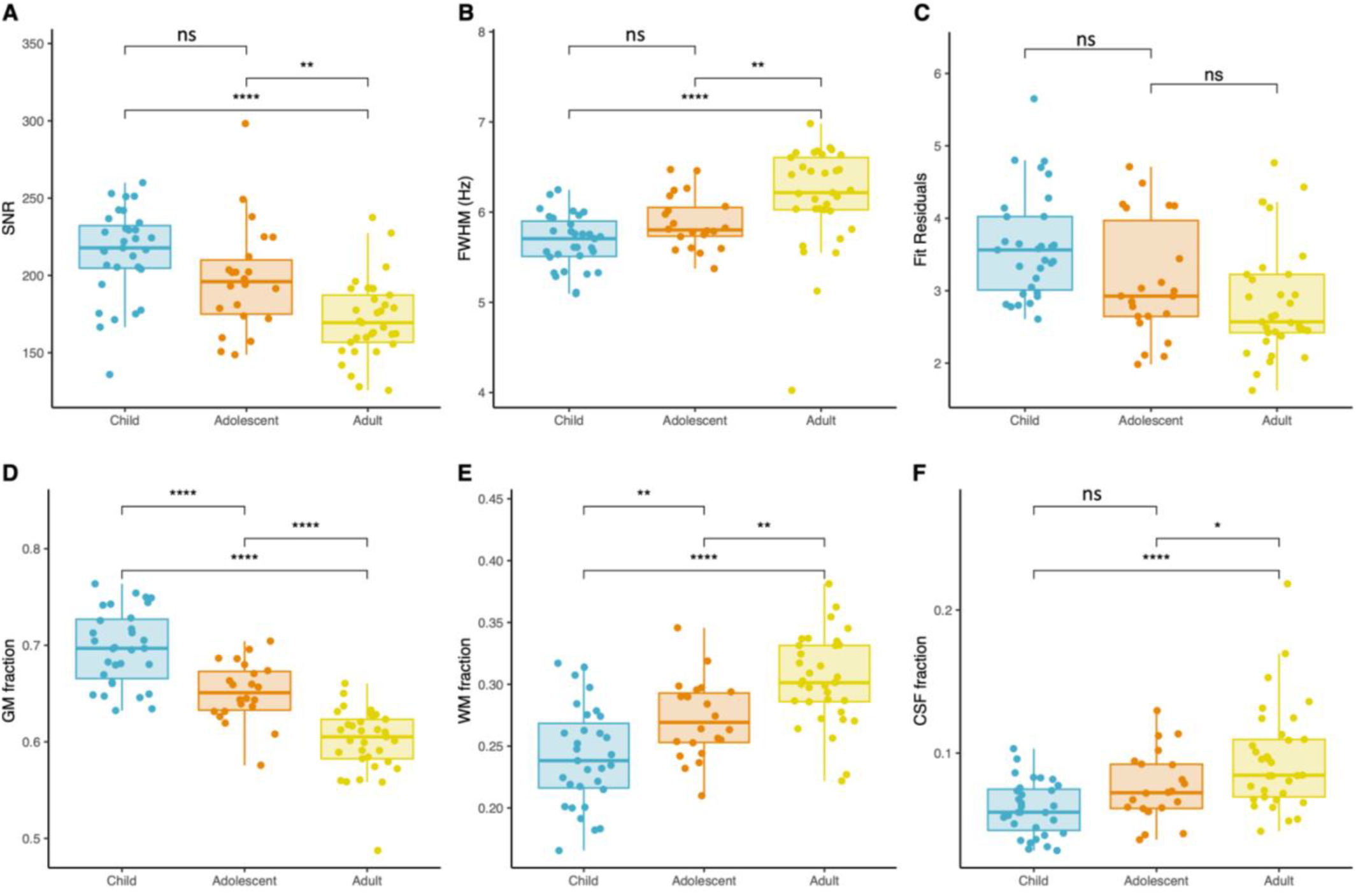
Quality metric differences across age groups. Following ANCOVA, Tukey post-hoc correction was used to find significant differences in QM between the age groups for (A) SNR, (B) FWHM, (C) fit residuals (OFF-ON), (D) voxel GM fraction, (E) voxel WM fraction, (F) voxel CSF fraction. ns: non-significant, *p < 0.05, **p < 0.005, ***p < 0.001, ****p < 0.0001.

### Tissue fractions

Age category showed a significant main effect on voxel tissue fractions (GM, WM, CSF; **Table 2**). Voxel GM fraction significantly decreased between age groups from children to adolescents to adults (**Figure 2**), and negatively correlated with age (Pearson’s r = −0.78, p < 0.0001). Voxel white matter and CSF significantly increased between age groups from children to adolescents to adults (**Figure 2**), positively correlating with age (Pearson’s r = 0.65, p < 0.0001 and Pearson’s r = 0.42, p < 0.0001 respectively). Note that data presented below as estimated metabolite concentrations in institutional units (i.u) is corrected for differences in voxel tissue composition and is consistent with creatine-referenced data as internal validation.

### Neurochemistry per age group

Mean estimated metabolite concentrations (i.u) and metabolite creatine ratios (/tCr) are reported for each age group in supplementary table 1; **Figure 3** (N = 86; 31 children, 22 adolescents and 33 adults). Age category showed a significant main effect on estimated concentrations (i.u) of tCr, tCho, Glx, and Glu (residuals of fit, sex and IQ as covariates; p < 0.05; see supplementary table 1). For creatine-ratio metabolite concentrations (/tCr), age category showed a significant main effect on tNAA, tCho, Glx, Glu and GABA+ (see supplementary table 1).

**Figure 3.**
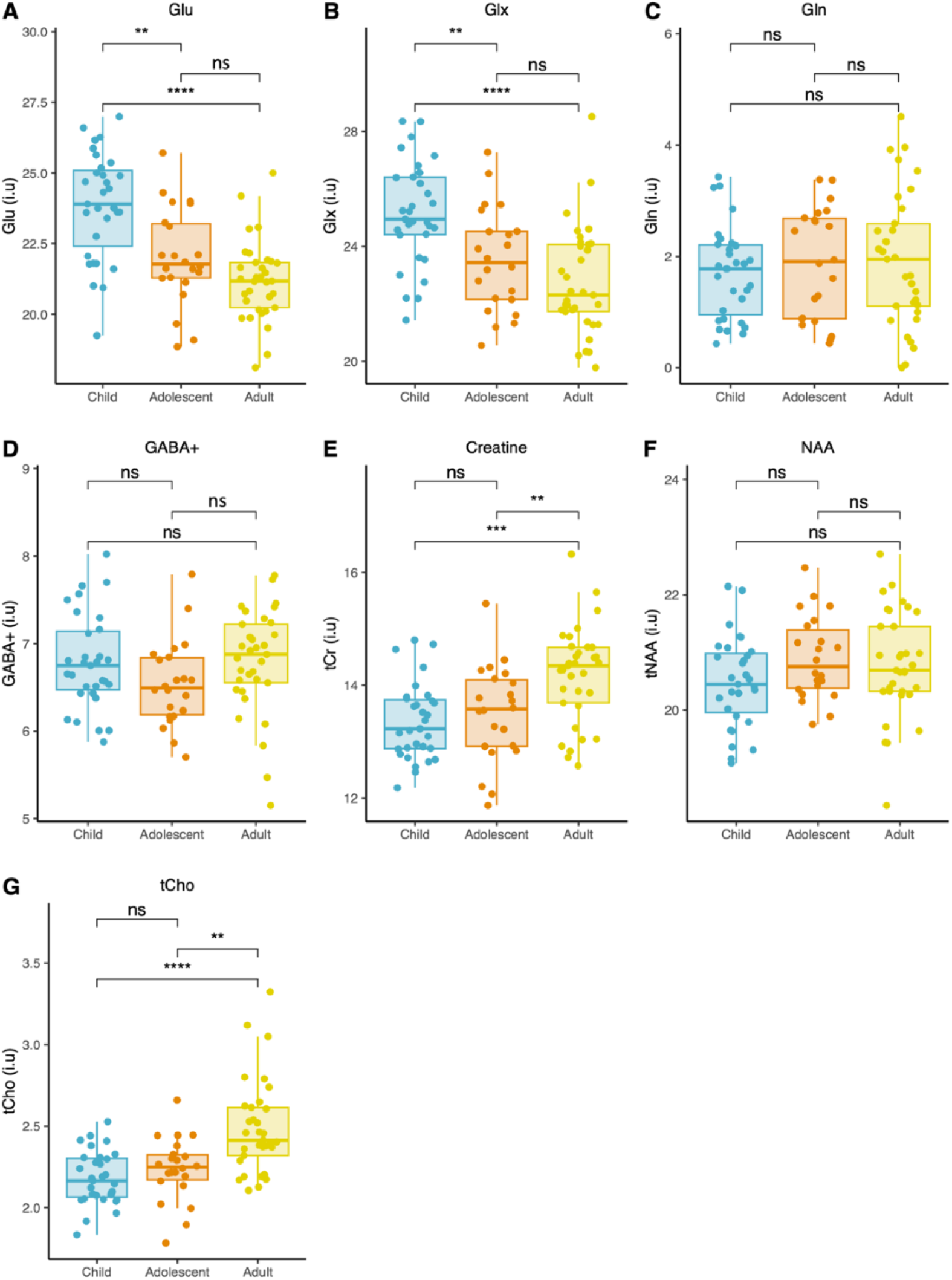
(A-G) Tukey post-hoc correction was used to isolate significant differences in estimated metabolite concentrations (i.u) between age groups. Significant differences are indicated via connecting lines. ns: non-significant, *p < 0.05, **p < 0.005, ***p < 0.001, ****p < 0.0001.

Sex had a significant main effect on estimated tCho and Gln only (F_2,83_ = 9.183, p < 0.0001; F_2,83_ = 5.154, p <0.05 respectively). Wilcoxon signed rank tests were used to assess differences in Gln and tCho concentrations between sexes per age group. After multiple comparison corrections (Bonferroni) only adult males had significantly greater concentrations compared to adult females of estimated tCho (2.35 (0.17) versus 2.60 (0.32); p <0.05). Significant differences in Gln concentrations between sex’s did not survive multiple comparison correction. The same was observed for metabolite creatine ratios (supplementary table 2, 3 & 4).

Fit residuals had a significant main effect on estimated tCho (F_2,83_ = 15.71, p < 0.05), Glx (F_2,83_ = 24.00, p < 0.001) and Glu concentrations (F_2,83_ = 34.72, p < 0.001). IQ had no significant main effect for any metabolite. The same was observed for metabolite creatine ratios (supplementary table 2).

ANCOVA Tukey post-hoc corrections were used to observe specific differences in mean metabolite concentrations between age groups, shown in **Figure 3**. Estimated concentrations (i.u.) of Glu and Glx significantly decreased between age groups from childhood to adulthood, while estimated concentrations of tCr and tCho significantly increased from childhood and adolescence to adulthood, further motivating our reporting and interpretation to focus on estimated concentrations (i.u). Gln, GABA+ and tNAA concentrations did not significantly differ between age groups after multiple comparison corrections. Creatine ratios of Glx, Glu, tNAA and GABA+ significantly decreased between age groups from childhood to adulthood (**Figure 4**). Mean creatine-ratio Gln did not significantly differ between age groups, while the creatine-ratio of tCho significantly increased from childhood to adulthood.

**Figure 4.**
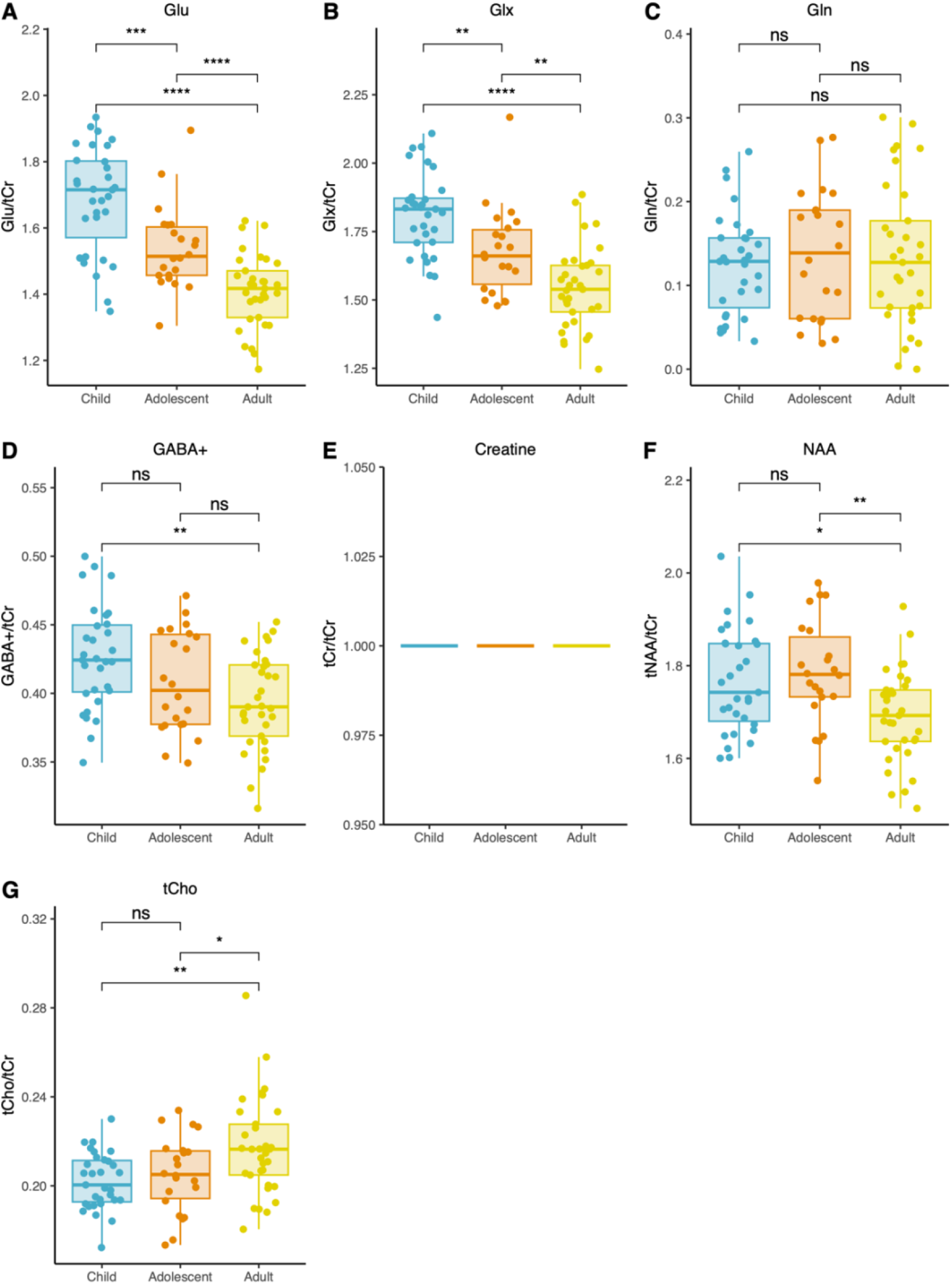
(A-G) Tukey post-hoc correction was used to isolate significant differences in creatine-ratio metabolite concentrations between age groups. Significant differences are indicated via connecting lines. ns: non-significant, *p < 0.05, **p < 0.005, ***p < 0.001, ****p < 0.0001.

### Linear effects of age per age group

To assess linear associations of metabolite concentrations with age per age group, the Pearson correlation coefficient was calculated for each metabolite per age group. Significant correlations were isolated; for estimated Glu a negative correlation with age was observed in children (r_children_ = −0.5, p < 0.001). For estimated GABA+, a significant negative correlation between concentration and age was observed in children (r_children_ = 0.5, p < 0.005). For estimated tCr a significant positive correlation between concentration and age was observed in children (r_children_ = 0.37, p < 0.005). For estimated tNAA, a negative correlation between age and concentration was observed in adults (r_adult_ = −0.34, p = 0.0557), just missing significance. Similar results were observed for creatine-ratio metabolite data.

### Developmental modeling

#### Linear regression

Linear models with metabolite concentration as the dependent variable and age as the independent variable with sex, IQ, and residuals of fit as predictors are shown in **Figure 5**. The linear age effects were significant and negative for estimated concentrations of Glu (beta = −0.11, p < 0.001) and Glx (beta = −0.11, p < 0.001), and significant and positive for estimated concentrations of tCr (beta = 0.056, p < 0.001) and tCho, for males and females (beta_pooled_ = 0.022, p_pooled_ < 0.001, beta_males_ = 0.023, p_males_ < 0.001, beta_females_ = 0.017, p_females_ < 0.005). No significant linear age effects were observed for sex pooled or sex separated Gln concentrations (estimated and creatine ratios) and estimated concentrations of tNAA and GABA+. For creatine ratios (**Figure 5**), linear age effects were significant and negative for tNAA (beta = −0.0075, p < 0.001), Glx (beta = −0.013, p < 0.001), Glu (beta =-0.018, p < 0.001) and GABA+ (beta = −0.0020, p < 0.005) and significant and positive for tCho sex pooled and tCho males, but not females (beta_pooled_ = 0.0010, p_pooled_ < 0.005, beta_males_ = 0.0012, p_males_ < 0.005, beta_females_ = 0.000, p_females_ < 0.005).

**Figure 5.**
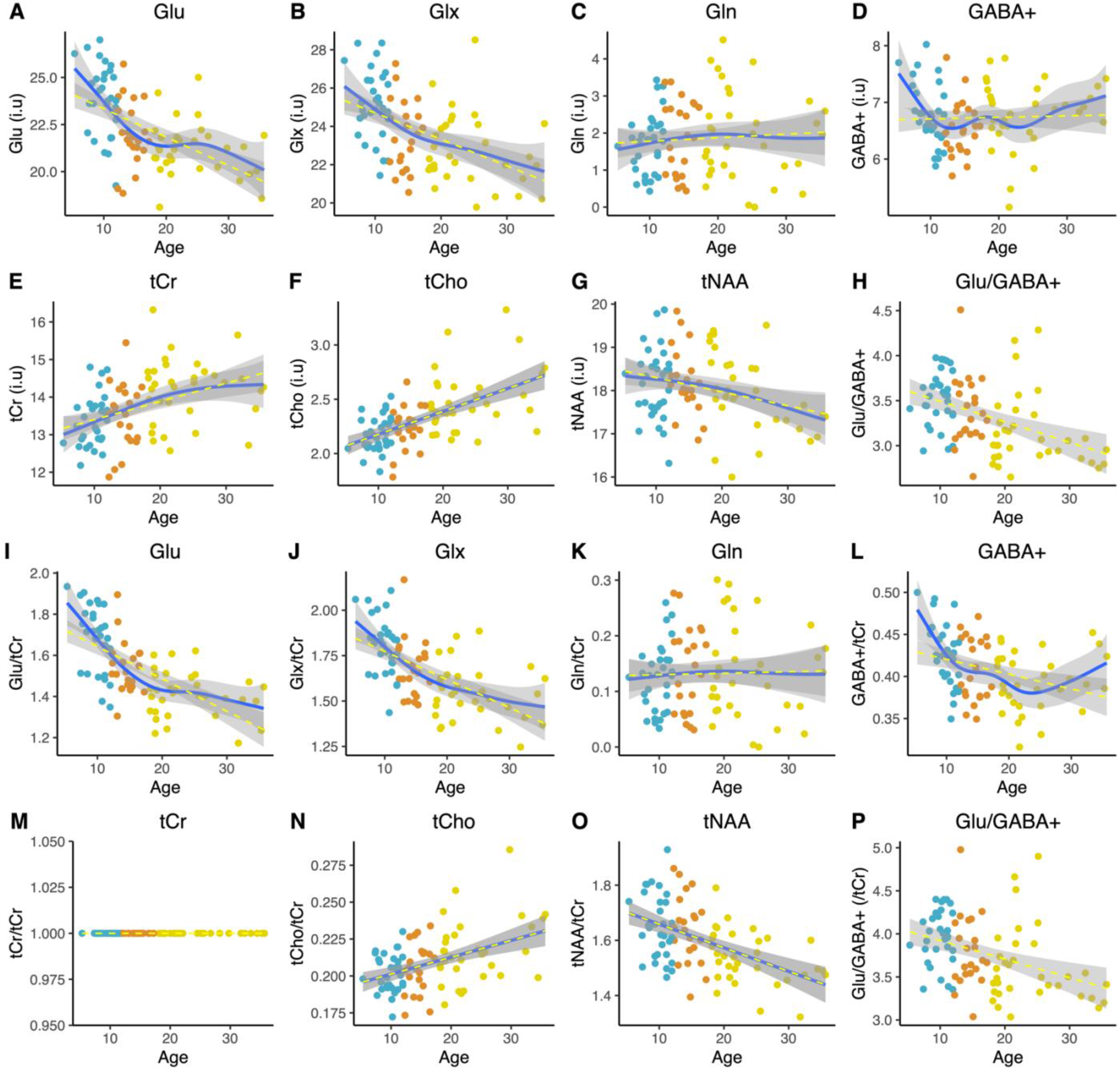
Linear (yellow) and GAM modelling (blue) of metabolite (A-G) estimated concentrations (i.u) and (I-P) creatine ratios over the lifespan. R^2^ values for GAMS and linear regression models are shown. Also shown are Pearson correlation coefficients (r) of Glu/GABA+ ratios for estimated (H) and creatine-ratio data (P) with age. Blue point = child, orange point = adolescent, yellow point = adult.

#### General Additive Models

GAMS are shown in **Figure 5**. Significant non-linear effects of age were identified for estimated concentrations of tCr (edf = 2.03, p < 0.005), Glu (edf = 2.954, p < 0.001), Glx (edf = 1.263, p < 0.001), tNAA (edf = 1.035, p < 0.001) and GABA+ (edf = 4.223, p < 0.05). Significant linear effects of age were observed for tCr (edf = 1, p < 0.001). As to the shape of the GAMs, **Figure 5** shows that estimated concentrations (i.u) of Glu, Glx and GABA+ decline relatively sharply in childhood and adolescence. Glx and Glu then more gradually decrease in early adulthood. tNAA concentrations (i.u) gradually decline from early adulthood. Concentrations of tCr (i.u) increase steeply from childhood to early adulthood, while tCho concentrations (i.u) increase linearly throughout the lifespan, from childhood to adulthood. For Gln (sex pooled; i.u) we observe no clear trends / changes in concentration across the lifespan.

Creatine-ratio concentrations show very similar trajectories to estimated concentrations for Glu, Glx, Gln, tCho and tNAA (**Figure 5**). GABA+/tCr however differs by decreasing across adolescence and adulthood (estimated concentrations remain relatively stable, with no net decrease).

Sex separated trajectories for Gln and tCho are found in supplementary figure 1. For Gln non-linear trajectories (GAM) were not significant for both sexs. For tCho male and female trajectories are similar, with gradual increases tCho concentrations across the lifespan, this trend was more linear in females than males (edf_females_= 1, p < 0.05, edf_males_ = 1.388, p < 0.05)

Finally, a significant negative correlation with age (across the whole dataset) was observed for estimated Glx:GABA+ ratios (r = −0.34, p < 0.005) and Glu:GABA+ (r = −0.42, p < 0.001; **Figure 5**). This was also observed for metabolite creatine ratios.

### Metabolite interactions

Correlations between neurometabolite concentrations differ between age groups. We were particularly interested in GABA and glutamate metabolism and thus, GABA+, Glu, Gln and Glx correlations are shown in **Figure 6** (see supplementary figure 2 for all other metabolite pairs). In childhood (r = 0.7; p < 0.001), adolescence (r = 0.61, p < 0.05) and adulthood (r = 0.35, p < 0.05) GABA+ significantly positively correlates with glutamate. Glx and GABA+ significantly positively correlate in children (r = 0.49, p < 0.005) and adolescents (r = 0.46, p < 0.05). Gln and GABA+ significantly negatively correlate in children only (r = −0.47, p < 0.05). In all age groups Glx and Glu significantly positively correlated (p < 0.0001).

**Figure 6.**
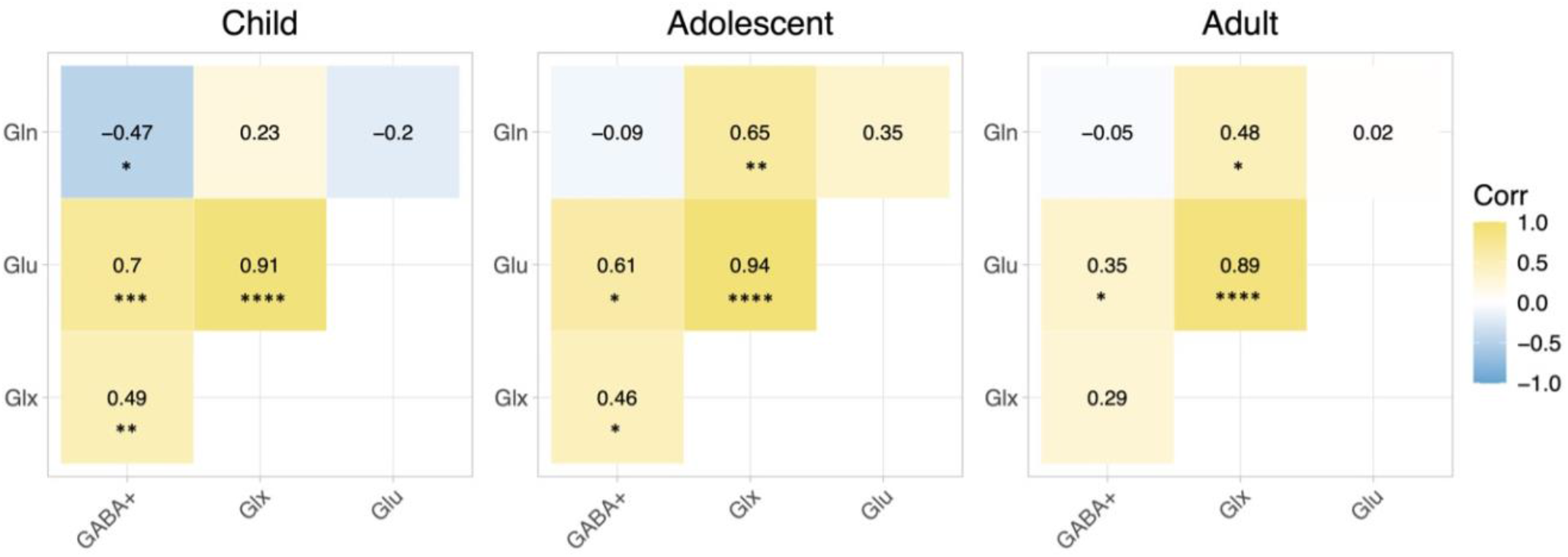
Correlation matrices for child, adolescent and adult estimated metabolite concentrations (i.u), with the Pearson correlation coefficients shown. Yellow = positive correlation, blue = negative correlation and white = no correlation. Significant correlations are shown *p < 0.05, **p < 0.005, ***p < 0.001, ****p < 0.0001.

### Correlations with cognitive function

Concentrations of GABA+ (i.u) significantly positively correlated with recognition memory (r = 0.27, p = 0.0126; **Figure 7**). The linear regression model (recognition memory ∼ age + GABA+) was used to adjust for the effect of age. When age was held constant, GABA+ concentrations (i.u) still showed a significant positive linear relationship with recognition memory scores (beta_pooled_ = 0.100, p_pooled_ < 0.01122; Figure 7C) across all ages. For Glu/Glx (i.u), we observed no significant relationship with recognition memory scores (supplementary table 5). Similar results were observed for creatine ratio data.

**Figure 7.**
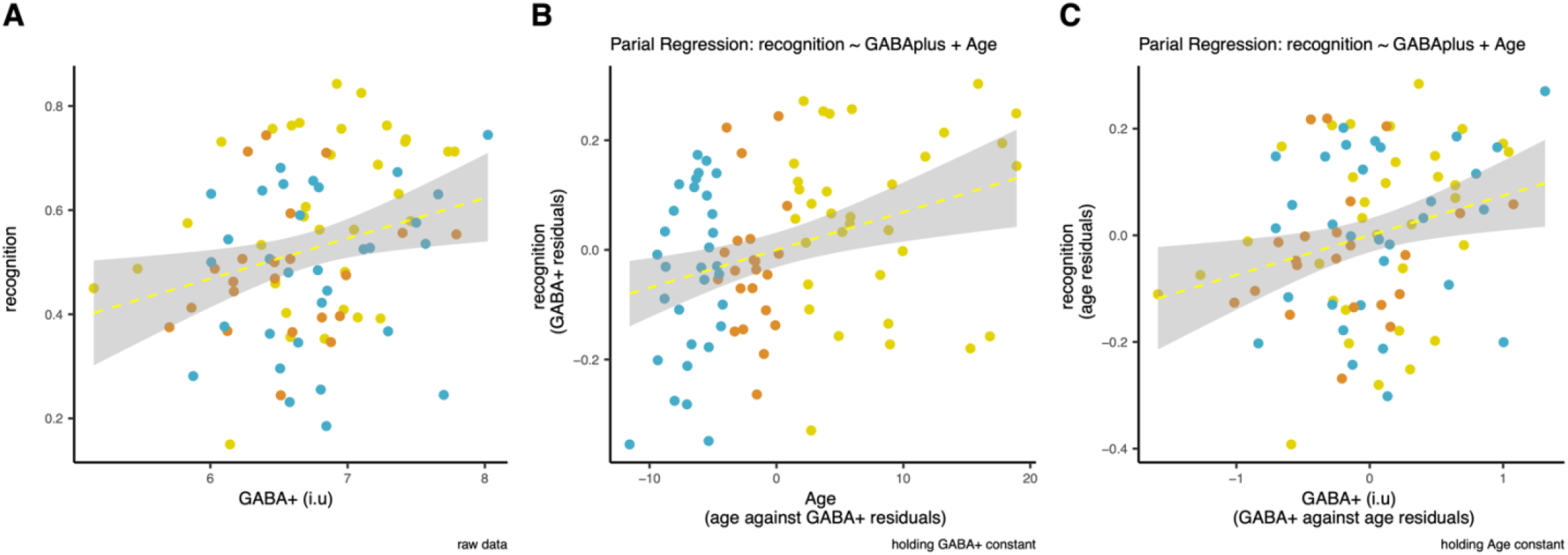
(A-B) Across all age groups, GABA+ concentrations (i.u) positively correlate with recognition memory scores (r = 0.27, p < 0.05). (B) Partial regression plot for the relationship between recognition memory scores and age when GABA+ (i.u) is held constant. (C) Partial regression plot for the relationship between recognition memory and GABA+ (i.u) when age is held constant. * indicates significance at p < 0.05. Grey bars indicate standard error. Blue point = child, orange point = adolescent, yellow point = adult.

## 4. Discussion

### Developmental trajectories

We used edited ^1^H-MRS to determine the lifespan trajectory of several neurometabolites, and their association with individual differences in cognition, in a large sample from childhood to early adulthood. Our primary data shows that the trajectories of metabolite concentrations are not strictly linear, but rather asymptotic. For example, estimated Glu concentrations show a sharp decrease in late childhood transitioning into early adolescence, before tapering off less steeply into early adulthood. Furthermore, we show that differences in cognitive function across development can be explained, in part, by differences in neurometabolism, specifically that of GABA+, the main inhibitory neurotransmitter. As such, the characterised developmental changes in neurometabolites are potentially (mechanistically) associated with functional development and so may represent valid therapeutic targets for atypical neurodevelopment whereby cognitive processes are disrupted. Below, we briefly discuss the relevance of each metabolite, as well as the limitations to our findings.

#### Glutamate

Our findings show, for both estimated and creatine-ratio Glu and Glx concentrations, a sharp (and significant) decrease from early childhood to adolescence. MRS measures of Glu are thought to be largely localised to the presynaptic terminal of glutamatergic neurons (Posse et. 2007; Sailasuata et al. 2008; Takado et al. 2019). A sharp decrease in Glu concentrations (i.u) across childhood and adolescence may thus reflect developmental pruning and refinement of excitatory synapses in the grey matter of our voxel (O’Leary et al. 1981; Huttenlocher and Dabholkar 1997; Selemon et al. 2013).

Decreases in neuronal/glial synthesis of Glu from Gln may also contribute, as studies in rats find that the cerebral expression of glutaminase, the enzyme that converts Gln to Glu, decreases from postnatal day 14 through to adulthood (Boulland et al. 2003; Shimizu et al. 2017;). Our Gln trajectories however do not support this, as we observe no significant changes in Gln concentrations (i.u or creatine ratios) across development, although this might be due to technical reasons (see below). Childhood decreases in Glu/Glx concentrations are consistent with observations in other MRS studies both within the parietal lobe (Volk et al. 2019), and in in other brain regions (the basal ganglia, and occipital lobe; Ghiselli et al. 2015; Shimizu et al. 2017).

Imaging studies have demonstrated that brain growth and development slows in early adulthood as the brain reaches maturity (Bethlehem et al., 2022). We indeed observe more gradual decreases in Glx/Glu concentrations at this time (i.u and creatine ratios). Studies observing older cohorts find a significant negative correlation between Glu concentrations and age in multiple brain regions including; parietal lobe (Gao et al., 2013, 20 - 76 years); anterior cingulate cortex (Hädel et al. 2013: Glu, 19 - 55 years), motor cortex (Kaiser et al. 2005: Glu, 24 - 68 years), frontal lobe (Sailasuta et al. 2008: Glu, 21 - 71 years; Gao et al. 2013: Glx, 20-76 years; Marsman et al. 2013: Glu, 18-31 years), basal ganglia, striatum (Sailasuta et al. 2008: Glu, 21 - 71 years; Ghisleni et al. 2015: Glu, 8 - 25 years). As such PPC Glu concentrations likely continue to decrease out of the scope of this study across mid/late adulthood. Of note, Sailasuta et al. 2008 (21 - 71 years) observed significant decreases in parietal Glx in adult men specifically, however, in a younger, larger cohort we observed no effect of sex on Glx/Glu trajectories.

The poor spectral separation achieved at the field strength used (3T) means it can be challenging to quantify individual concentrations of Glu and Gln (Zhang et al. 2016). As such, for transparency, we report both Glx and Glu. Importantly however, our measure of Glu-only was obtained through linear combination fitting, whereby it is assumed that the majority of Glx is indeed Glu. This may contribute to the observed significant positive correlation between estimated Glx and Glu concentrations in all age groups and the absence of a correlation between estimated Glx and Gln. Our failure to isolate significant changes in Gln concentrations may thus also be an artefact of this fitting constraint.

#### GABA

We found that GABA+ concentrations (estimated and creatine ratios) have a non-linear relationship with age, showing a relatively sharp decline across childhood before plateauing in adulthood. This non-linear approach better explains the variance of GABA+ concentrations over age compared to linear models (greater R^2^), while the significant negative corelation between GABA+ and age in children (consistent with Ghisleni et al. 2015), which is absent in adolescents and adults, is consistent with this model. We should note that these results conflict with a recent meta-analysis by Porges et al. 2021 and findings by Saleh et al. 2020, which described an increase in frontal GABA+ concentrations during late childhood (8 - 12 years). These differences are likely explained by voxel placement, as biochemical development of the PPC differs markedly in timing to the frontal lobe, which is one of the last brain regions to mature (Ouyang et al. 2019). Increases in GABA+ concentrations (thought to reflect myelination and increased synaptic activity; Porges et al. 2021) may thus occur earlier than the scope of this study in PPC. Similar increases in Glu may also precede our trajectories, as Perdue et al. 2022 observed that temporal-parietal Glx concentrations increased between 2 - 6 years of age before decreasing from mid childhood, as we observed in the PPC in our data.

A hallmark of postnatal neurodevelopment is maturation of the GABAergic system, characterised as a switch from excitatory to inhibitory neuronal responses triggered by GABA’s bindings to GABA receptors (Dzhala et al. 2005), developmental changes in GABA tonic and presynaptic receptor currents (Fritschy et al. 1994) and changes to GABA receptor subunits (Okada et al. 2000). MRS GABA+ signal is thought to reflect the degree a circuit can employ this GABAergic inhibition, mainly composed of tonically active (extra-cellular) GABA (as opposed to synaptic GABA which mediates phasic inhibition; Rae et al. 2014; Puts et al. 2017; Hermans et al. 2018). As such, childhood decreases in GABA+ concentration (concurrent with decreases in Glu) likely reflect this post-natal GABAergic maturation, and the establishment of a homeostatic balance between the newfound inhibitory activity of GABA and the excitatory activity of glutamate, essential for typical brain functioning and development (Rubenstein 2010; Shimizu et al. 2017).

We observe this balance changing over the course of development as Glu:GABA+ ratio significantly negatively correlates with age, in keeping with an increasing shift in the balance of inhibitory/excitatory brain activity towards inhibition with age (Ghisleni et al. 2015). This is consistent with several observations of a developmental decrease in cortical excitatory activity but not inhibitory activity in mice (amplitude of excitatory post synaptic potentials and release probability; Reyes and Sakmann 1999; Oswald and Reyes 2008; Feldmeyer and Radnikow, 2009; Zhang et al. 2011). Accordingly, our results show that estimated GABA+ concentration remain relatively stable in early adulthood/adolescence while, while Glu concentrations continue to gradually decrease in early adulthood and adolescence, likely due to this developmental depression of excitatory activity only. Increased neuronal excitability (greater Glu:GABA+ ratio) in early development is thought to facilitate a critical period of synaptic plasticity, essential for the early establishment and refinement of neural circuitry before increasing inhibition/reduced excitation (and so lower Glu:GABA+ ratio) with local circuit maturation (Citri and Malenka 2008; Zhang et al. 2011).

#### Choline

In contrast to GABA+/Glu, we find a significant increases in tCho and tCr concentrations across childhood to early adulthood (estimated and creatine ratios). For tCho, this relationship appears to be strictly linear (edf = 1) within the age range studied.

Furthermore, sex interactions were observed; a steeper increase in tCho concentrations across the lifespan in males compared to females, as adult males had significantly greater concentrations of PPC tCho compared to adult females. The potentiating effects of oestrogen on choline acetyltransferase expression, an enzyme responsible for tCho conversion into acetylcholine, may underpin these sex differences for tCho trajectories (Bohacek et al. 2008; Hädel, et al. 2013). This may also explain why we observe sex differences only *after* puberty, with no significant difference in tCho concentration observed in children.

Age-associated increases in tCho concentrations have been observed in a number of brain regions in adult only cohorts (Chang et al. 1996 (19 - 78 years), Lind et al. 2020 (ACC; 18 - 79 years), Ding et al. 2016 (WM; 20 - 70 years), Schmitz et al. 2018 (22 - 73 years; whole brain), Pfefferbaun et al. 1999 (frontal lobe; 25-73 years old), Schmitz et al. 2019 (whole brain MRSI; 22 - 73 years); Sailasuta et al. 2008 in (frontal WM;21 - 73 years). We observe this relationship to be true in childhood also. As a marker of neuron integrity, as well as myelination (Rae et al. 2014) increasing choline concentrations may represent developmental increases in PCC structural connectivity from childhood to adulthood (Supekar et al. 2010; Sato et al. 2014; Fan et al. 2021; Fair et al. 2008). In part this is underpinned by the increases in nerve fibre density and myelination, consistent with our observations that white matter voxel fractions increase with age (Fair et al. 2008; Sato et al. 2014; Fan et al. 2021).

#### Creatine

For tCr (i.u) we observed no significant sex interactions, however non-linear modelling found a significant quadratic interaction of tCr with age (edf = 2); ‘steeper’ increases in concentrations (i.u) during adolescence before levelling off in adulthood (the positive correlation of creatine concentrations with age in childhood lost in adulthood). Age-associated increases in tCr concentrations have again been observed in several brain regions in narrower age ranges; (Blüml et al. 2013; parietal lobe, Marsman et al. 2013; medial frontal cortex; 18 - 31), Schmitz et al. 2018 (whole brain MRSI; 30 - 67 years), Blüml et al. 2013 (cerebral GM; 5 months - 12 years) and Kaiser et al. 2005 (motor cortex and corona radiata; 24 - 68 years). We however show (through non-linear modelling) that this increase to the most significant and steeper during childhood and adolescence. This likely reflects the changing metabolic activity of neurons and glial cells during this period of great neurodevelopmental change.

#### tNAA

For tNAA (i.u) we observe a significant non-linear interaction with age, with a gradual decline in concentrations from early adulthood. This is consistent with age related declines in tNAA/NAA concentrations previously described in adult cohorts only (Saleh et al. 2019; occipital lobe; 18 - 87 years), Kirov et al. 2021 (20 - 90 years; whole brain MRSI), Ding et al. 2016, Schmitz et al. 2018 (20-70 years; whole brain MRSI) and Kasier et al. 2005 (24 - 68 years; motor cortex). Such adulthood tNAA declines may reflect a slow and increasing mitochondrial impairment due to the gradual accumulation of oxidative stress (reactive oxygen species), leading to increasing mitochondrial DNA mutations as the brain begins to age (Zuendorf et al. 2003; Demougeot et al. 2001; De Moura et al. 2010; Chistiakov et al. 2014; Siddiqui et al. 2016). Subsequently, NAA synthesis in the neuronal and glial mitochondria undergoes a gradual decline (Patel and Clark 1979; Rae et al. 2014; Siddiqui et al. 2016).

It is important to note that whilst mostly consistent, we did observe differences in metabolite trajectories between estimated concentrations and creatine ratios. For example, creatine-ratio parietal GABA and tNAA concentrations showed a significant negative linear association with age, while estimated GABA+ and tNAA concentrations had a non-linear association only (see above). The observed age-related increases in creatine concentrations likely drive these differences and so creatine-ratio trajectories may fail to reflect real changes in tissue physiology. This reinforces the need to report both estimated (i.u) and creatine-ratio concentrations (and preferably estimated concentrations with the appropriate corrections) and be transparent in reporting (also see Lin et al. 2021).

In summary, the characterised metabolite trajectories are likely underpinned by the developmental refinement, maturation, and myelination of excitatory and inhibitory brain circuitry in the postnatal period. Trajectories diverge due to the differing developmental process(s) governing each metabolite, spanning from mitochondrial activity and neuronal energy consumption to synaptic pruning.

### Metabolite-metabolite interactions

GABA+ and Glu were found to significantly positively correlate in all age groups, likely reflective of their shared metabolite pathway; GABA is synthesised in inhibitory neurons by the decarboxylation of Glu via the enzyme glutamic acid decarboxylase (GAD); see review Martin and Tobin 2000. As such, their concentrations are biologically linked, and thus it is encouraging that we observe this in our MRS measures. In the brain, Gln is synthesised in astrocytes, released, and taken up by neurons for GABA and Glu synthesis (Bak et al. 2006). These key differences in spatial localisation of GABA/Glu (neurons) and Gln (neurons and astrocytes) may contribute to the lack of linear interaction between GABA/Glu and Gln concentrations in our study, with likely more complex (non-linear) interactions between their concentrations across development as neuronal signalling pathways mature.

### Memory correlates

Across all age groups PCC GABA+ concentrations were significantly positively associated with recognition memory scores. We did not see any significant relationship between PPC Glx/Glu concentrations and memory scores, supporting the specificity of this association.

PPC GABA may contribute to recognition memory performance via the tuning of pyramidal neuron synchronisation within the hippocampal-parietal network. The PPC, acting within a hippocampal-parietal memory network, has been increasingly implicated in episodic memory (including recognition; Donalson and Rugg 1998; Henson et al. 1999; Wagner et al. 2005; Cabeza et al. 2008; Vincent et al. 2006; Ciaramelli et al. 2008; Wang et al. 2010; Riggins et al. 2016; Ngo et al. 2017). Within this hippocampal-parietal network, theta (low frequency) and gamma (high frequency) oscillations of pyramidal neuron activity are also associated with episodic memory performance (Uhlhaas et al. 2010; Nyhus and Curran 2011; Heusser et al. 2016; Natu et al. 2019; Griffiths et al. 2021). GABA has been shown to regulate this oscillatory activity (gamma; Cobb et al. 1995; Whittington et al. 1995; Traub et al. 1996; Chen et al. 2014). This evidence thus provides a link between GABA concentrations and recognition (a type of episodic) memory performance.

### Limitations

We note several limitations to our study. First, metabolite concentrations across all ages in our study population were post-hoc corrected using literature derived (adult) tissue water and metabolite T1 and T2 relaxations values (Gasparovic et al. 2006; Oeltzschner et al. 2020; Hui et al. 2022). Due to changes in brain tissue composition and particularly water content, these values likely vary with age (Kirov et al., 2008; Christiansen et al. 1993). Furthermore, macromolecule (MM) contributions to the GABA+ signal were also post-hoc corrected using existing MM basis set functions modelled on data from participants from 20 - 69 years old (Oeltzschner et al. 2020; Hui et al. 2021). While Hui et al. 2021 found no significant age associated changes in MM signals, it cannot be ruled out that childhood changes in MM contribution do not underpin the observed decreases in GABA+ concentrations (Bell et al. 2021). Macromolecule supressed editing can overcome this limitation, however, it is highly motion sensitive and achieves a significantly lower SNR compared to MM unsuppressed MEGA-PRESS (Mikkelsen et al. 2017; Bell et al. 2021).

Beyond T1 and T2 relaxation and MM contributions, basis sets are also tailored towards adult data and which may explain the increased fitting errors found in the younger cohort. However, we note that our quality metrics have low variability and are within normative ranges (Mikkelsen et al. 2017) and thus are unlikely to broadly affect our results. We additionally used fit residuals as a covariate in our linear and non-linear models, with age still significantly predicting metabolite concentrations and unchanged levels of significance. We also report very similar results for creatine-referenced metabolite levels, for which no post hoc corrections (T1/T2/tissue) are made, and as such it is unlikely that the trends identified in water referenced data are artefacts of post-hoc processing. In the future, characterisation of metabolite and tissue water T1/T2 relaxation and MM signal contributions across the lifespan (and for specific brain regions) will enable more tailored corrections of metabolite concentrations better addressing these accuracy concerns.

Of note, within our cohort, we observed IQ scores to be significantly greater in adults compared to children and adolescents. As such, IQ was included as a covariate in our linear and non-linear models. Despite this, age still significantly predicted metabolite concentrations and levels of significance did not change. We also identified no significant effect of IQ on metabolite concentrations, when added to ANCOVA’s, linear regression and non-linear models. Thus, we are confident that this has not had a broad effect on our results overall.

Finally, as a cross-sectional study, individual variation is likely to contribute to theorised trajectories, especially past 30 years of age where the data was sparser and so non-linear models became more heavily influenced by individual data points. Furthermore, while adjusted R^2^ values obtained from GAM models were greater than adjusted R^2^ values obtained by linear regression for all metabolites (**Figure 5**), this may partially stem from the nature of the fitting method. In fact, we report linear regression models to prevent overinterpretation of non-linear data trends that maybe artefacts of GAM overfitting. In future, larger (longitudinal) datasets must be analysed, increasing power and reliability of identified data trends.

## Conclusion

We characterised the lifespan trajectories of 6 essential MRS metabolites a large, single site dataset, measured from a PPC voxel. For GABA, Glx, tNAA and tCr we observed significant non-linear trajectories; Glx/Glu and GABA+ concentrations steeply (and significantly) decreased across childhood before tapering off into early adulthood, perhaps reflective of the fine-tuning of excitatory and inhibitory tone during early developmental circuit refinement. tNAA concentrations displayed a gradual non-linear decrease from early adulthood, while tCr concentrations increased in childhood and adolescence. tCho has a linear and positive association with age. For Gln we observed no age-related differences in concentration. Finally, our characterised trajectories are associated with cognitive outcomes as GABA+ concentrations significantly positively correlate with recognition memory scores across post-natal development, which may reflect the role of PPC GABAergic activity in modulating the developing hippocampal-parietal memory network.

## 5. Funding

This work was supported by R01 EB032788, R01 EB016089, R01 EB023963, P41 EB031771 and S10 OD021648, and by the Johns Hopkins Therapeutic Cognitive Neuroscience Fund (XC and RL; Grant Number 80026224). ART is supported through a Medical Research Council PhD award [MR/P502108/1]. TA is supported by an MRC Transition Support Award [MR/V036874/1]. TA and NP received support from the Medical Research Council Centre for Neurodevelopmental Disorders, King’s College London [MR/N026063/1]. XC is supported by the Natural Sciences and Engineering Research Council of Canada (XJC; RGPIN-2020-05520), Canada First Research Excellence Fund, awarded to McGill University for the Healthy Brains for Healthy Lives initiative, and the Canada Research Chairs program.

## 6. Data availability

Our analytical code is available through: [LINK].

Data are available through: [LINK].

Osprey 2.4.0 is available through: https://github.com/schorschinho/osprey

## Notes

### Competing Interest Statement

The authors have declared no competing interest.

