## supplementary materials for "The developmental trajectory of ^1^H-MRS brain metabolites from childhood to adulthood"

Supplementary table 1. Mean (SD) estimated (i.u) and creatine-ratio (/tCr) metabolite concentrations for each age group.

| Metabolite | Child | Adolescent | Adult | p value |
|---|---|---|---|---|
| tCr (i.u) | 13.358 (0.675) | 13.500 (0.858) | 14.181 (0.857) | 0.000532*** |
| tCr/tCr | - | - | - | - |
| tNAA (i.u) | 20.473 (0.784) | 20.901 (0.719) | 20.797 (0.915) | 0.138 |
| tNAA/tCr | 1.767 (0.112) | 1.786 (0.113) | 1.690 (0.099) | 0.00163** |
| tCho (i.u) | 2.180 (0.163) | 2.236 (0.195) | 2.489 (0.287) | 1.55e-05**** |
| tCho/tCr | 0.202 (0.013) | 0.205 (0.017) | 0.217 (0.022) | 0.00224** |
| Glx (i.u) | 25.148 (1.782) | 23.522 (1.804) | 22.753 (1.878) | 0.0015** |
| Glx/tCr | 1.809 (0.156) | 1.676 (0.159) | 1.544 (0.149) | 3.87e-06**** |
| Glu (i.u) | 23.860 (1.877) | 22.038 (1.710) | 21.186 (1.405) | 8.62e-08**** |
| Glu/tcr | 1.684 (0.159) | 1.539 (0.126) | 1.409 (0.110) | 1.88e-08 **** |
| Gln (i.u) | 1.714 (0.822) | 1.872 (0.999) | 1.937 (1.195) | 0.649 |
| Gln/tCr | 0.125 (0.060) | 0.137 (0.077) | 0.134 (0.085) | 0.790 |
| GABA+ (i.u) | 6.787 (0.542) | 6.539 (0.489) | 6.794 (0.594) | 0.170 |
| GABA+/tCr | 0.426 (0.038) | 0.408 (0.037) | 0.394 (0.034) | 0.00135 ** |

Note: 31 children, 22 adolescents and 33 adults. p value indicates significance from multiple comparison ANCOVA's to test main effect of age group on metabolite concentrations (IQ, sex and residuals of fit as covariates). *p<0.05, **p<0.005, ***p<0.001, ****p<0.0001.

Supplementary Table 2. p values of intercepts obtained from multiple comparison ANCOVA's with estimated (i.u) or creatine ratio (/tCr) metabolite concentration as the dependant variable and IQ, sex, fit residuals and age group as independent variables.

| Metabolite | Estimated concentration (i.u) | | | | Creatine scaled concentration (/tCr) | | | |
|---|---|---|---|---|---|---|---|---|
| | IQ | sex | fit residual | age group | IQ | Sex | fit residual | age group |
| tCr | 0.220242 | 0.577032 | 0.128102 | 0.000532 *** | NA | NA | NA | NA |
| tNAA | 0.0658 | 0.1034 | 0.7428 | 0.1375 | 0.85506 | 0.11415 | 0.21797 | 0.00163 ** |
| tCho | 0.54334 | 0.00329 ** | 0.00063 *** | 1.55e-05 *** | 0.86082 | 0.00422 ** | 0.00491 ** | 0.02237 * |
| Glx | 0.4387 | 0.3143 | 5.32e-06 *** | 0.0015 ** | 0.210 | 0.649 | 1.45e-05 *** | 3.87e-06 *** |
| Glu | 0.463 | 0.730 | 2.91e-05 *** | 8.62e-08 *** | 0.202 | 0.466 | 2.50e-07 *** | 1.88e-08 *** |
| Gln | 0.8195 | 0.0259 * | 0.5973 | 0.6486 | 0.7262 | 0.0279 * | 0.7203 | 0.7895 |
| GABA+ | 0.681 | 0.966 | 0.318 | 0.170 | 0.50636 | 0.79732 | 0.87586 | 0.00135 ** |

Note: *p<0.05, **p<0.005, ***p<0.001, ****p<0.0001.

Supplementary table 3. Estimated (i.u) and creatine ratio (tCr) concentrations of tCho and Gln per age group per sex.

| Metabolite | Child | | Adolescent | | Adult | |
|---|---|---|---|---|---|---|
| | F | M | F | M | F | M |
| tCho (i.u) | 2.13 | 2.23 | 2.20 | 2.27 | 2.35 | 2.60 |
| | (0.12) | (0.19) | (0.19) | (0.20) | (0.17) | (0.32) |
| tCho/tCr | 0.20 | 0.20 | 0.20 | 0.21 | 0.20 | 0.23 |
| | (0.010) | (0.015) | (0.019) | (0.016) | (0.015) | (0.021) |
| Gln (i.u) | 1.59 | 1.83 | 1.57 | 2.12 | 1.54 | 2.27 |
| | (0.80) | (0.85) | (1.01) | (0.96) | (1.20) | (1.12) |
| Gln/tCr | 0.12 | 0.13 | 0.11 | 0.16 | 0.11 | 0.16 |
| | (0.059) | (0.062) | (0.077) | (0.075) | (0.083) | (0.082) |

Note: F = female, M = male.

Supplementary table 4. p values obtained from Wilcoxon Rank test for differences in estimated (i.u) and creatine ratio (/tCr) tCho and Gln concentrations between sexes for each age group.

| Metabolite | Child | Adolescent | Adult | All |
|---|---|---|---|---|
| tCho (i.u) | P = 0.0784 | P = 0.381 | P = 0.0106* | 0.0059 |
| tCho/tCr | P = 0.32 | P = 0.67 | P = 0.0021* | 0.0056 |
| Gln (i.u) | P = 0.626 | P = 0.254 | P = 0.0732 | 0.019 |
| Gln/tCr | P = 0.74 | P = 0.25 | P = 0.044 | 0.023 |

Note: * indicates significance after Bonferroni multiple comparison correction.

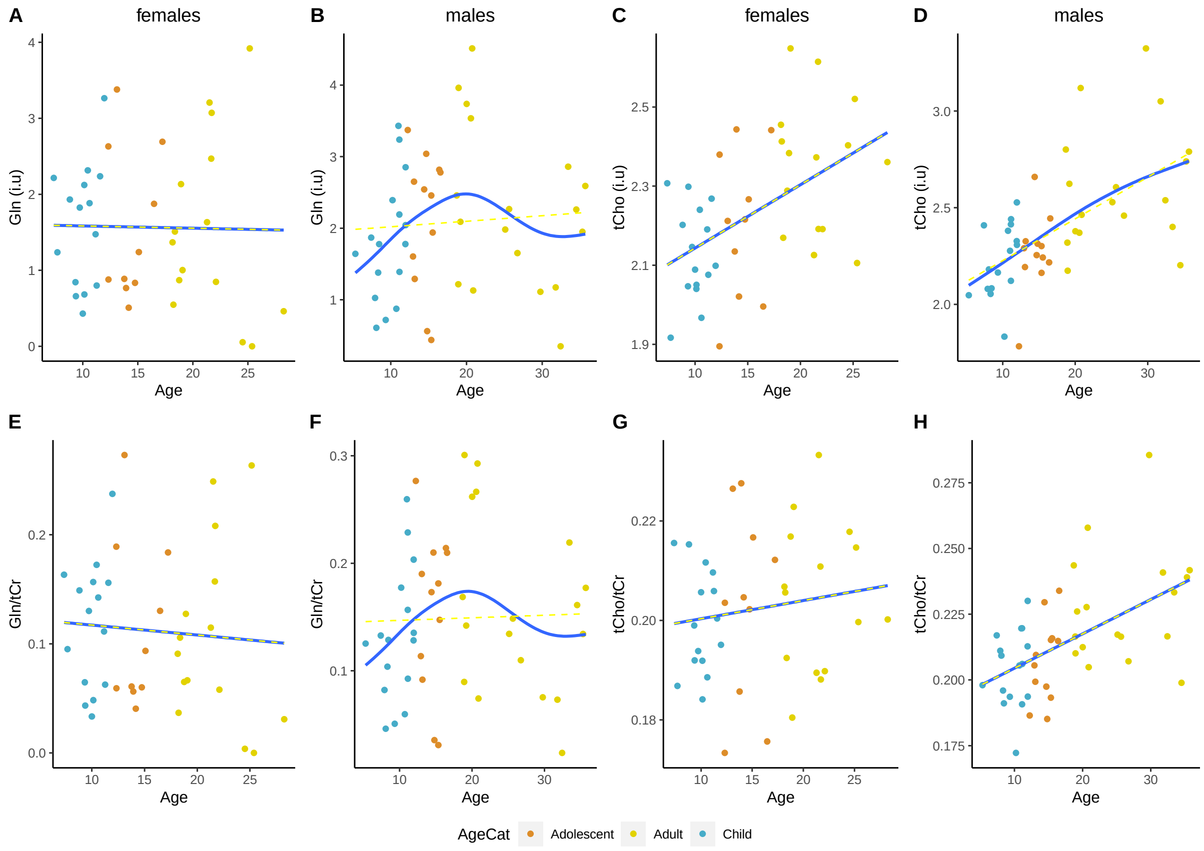


Supplementary figure 1. Linear (yellow) and non-linear GAM modelling (blue) of (A-D) estimated (i.u) and (E-H) creatine ratio (/tCr) metabolite concentrations of Gln and tCho across the lifespan when data was grouped by sex.

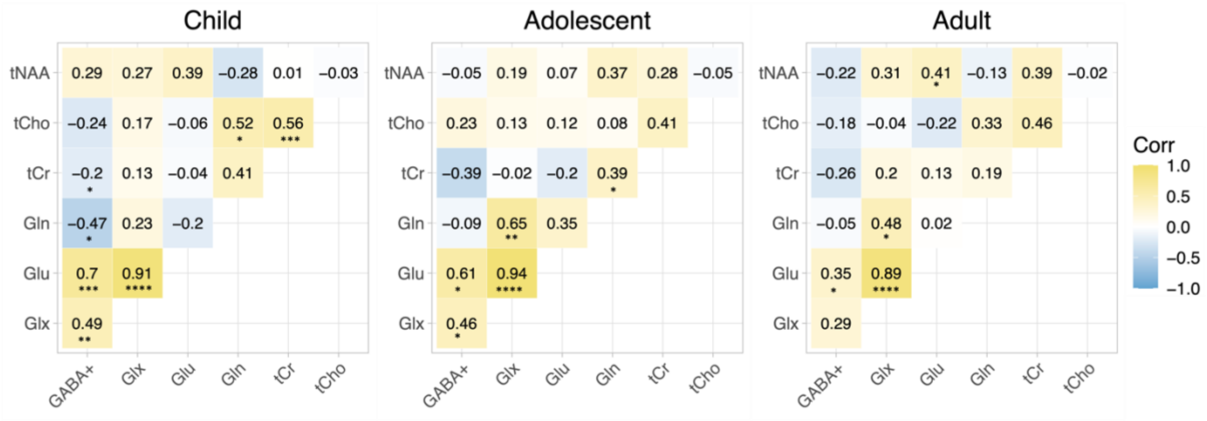


Supplementary figure 2. Correlation matrices for child, adolescent and adult estimated metabolite concentrations, with the Pearson correlation coefficients (r) shown. Yellow = positive correlation, blue = negative correlation and white = no correlation. Significant correlations are shown *p<0.05, **p<0.005, ***p<0.001, ****p<0.0001.

Supplementary table 5. Pearson correlation coefficients for estimated metabolite concentration and recognition, recollection and source memory scores.

| Metabolite | Pearsons r | p value |
|---|---|---|
| Glu: recognition | *-0.11* | *0.311* |
| Glu: recollection | *-0.18* | *0.0922* |
| Glu: source memory | *-0.17* | *0.121* |
| Glx: recognition | *-0.1* | *0.34* |
| Glx: recollection | *-0.16* | *0.147* |
| Glx: source memory | *-0.12* | *0.255* |
| GABA+: recognition | *0.27* | *0.0126** |
| GABA+: recollection | *0.19* | *0.0775* |
| GABA+: source memory | *0.18* | *0.105* |
| Age: recognition | *0.32* | *0.0026** |
| Age: recollection | *0.37* | *0.000405**** |
| Age: source memory | *0.31* | *0.00386 *** |

*Note:* *p<0.05, **p<0.005, ***p<0.001, ****p<0.0001.
